# Decoding Visual Spatial Attention Control

**DOI:** 10.1101/2023.08.05.552084

**Authors:** Sreenivasan Meyyappan, Abhijit Rajan, Qiang Yang, George R Mangun, Mingzhou Ding

## Abstract

In models of visual spatial attention control, it is commonly held that top-down control signals originate in the dorsal attention network, propagating to the visual cortex to modulate baseline neural activity and bias sensory processing. However, the precise distribution of these top-down influences across different levels of the visual hierarchy is debated. In addition, it is unclear whether these changes in baseline neural activity directly translate into improved performance. We analyzed attention-related baseline activity during the anticipatory period of a trial-by-trial voluntary spatial attention task, using two independent fMRI datasets, and two different analytic approaches. First, as in prior studies, univariate analysis showed that covert attention significantly enhanced baseline neural activity in higher-order visual areas contralateral to the attended visual hemifield, while effects in lower-order visual areas (e.g., V1) were weaker and more variable. Second, in contrast, multivariate pattern analysis (MVPA) revealed significant decoding of attention conditions across all visual cortical areas, with lower-order visual areas exhibiting higher decoding accuracies than higher-order areas. Third, decoding accuracy, rather than the magnitude of univariate activation, was a better predictor of a subject’s stimulus discrimination performance. Finally, the MVPA results were replicated across two experimental conditions, where the direction of spatial attention was either externally instructed by a cue or based on the participants free choice decision about where to attend. Together, these findings offer new insights into the extent of attentional biases in the visual hierarchy under top-down control, and how these biases influence both sensory processing and behavioral performance.

**Highlights:**

- Multivariate pattern analysis revealed the presence of top-down attentional biasing signals in all areas of the visual hierarchy whereas univariate analysis was not able to reveal the full extent of attentional biasing in the visual cortex.
- The decoding accuracy derived from the MVPA analysis but not the magnitude difference derived from the univariate analysis predicted the subject’s behavioral performance in stimulus discrimination.
- The MVPA results were consistent across two experimental conditions where the direction of spatial attention was driven either by external instructions or from purely internal decisions.

## Introduction

Extensive neurophysiological, neuroimaging, and lesion evidence suggests that frontoparietal cortical areas including the frontal eye field (FEF) and the intraparietal sulcus (IPS in humans) are the neural substrate underlying the control of voluntary attention (Astrand et al., 2015; Astrand et al., 2016; Bichot et al., 2015; Bisley & Goldberg, 2003; Bowling et al., 2020; Bressler et al., 2008; Buschman & Miller, 2007; Corbetta & Shulman, 2002; Gregoriou et al., 2009; Gunduz et al., 2012; Hopfinger et al., 2000; Ibos et al., 2013; Meehan et al., 2017). According to prevailing theory, in addition to maintaining the attention set, these areas also issue top-down signals to bias the responsiveness of sensory neurons in a goal-directed fashion in advance of target processing (Bressler et al., 2008; Carrasco, 2011; Chawla et al., 1999; Giesbrecht et al., 2006; Hopfinger et al., 2000; Kastner et al., 1999; Kastner & Ungerleider, 2000; Luck et al., 1997; Park & Serences, 2022; Snyder et al., 2021; Sylvester et al., 2007). One physiological sign of the influence of top-down control signals in visual cortex is the increase in baseline neural activity—baseline shifts—in visual cortex during anticipatory attention.

Attention-related baseline shifts have been observed during anticipatory attention in both humans and non-human primates. In single neuron recordings in nonhuman primates, Luck and colleagues (Luck et al., 1997) reported increased baseline neuronal firing rates in V2 and V4 (but not in V1) in neurons that encoded the to-be-attended information. In human electroencephalographic (EEG) and magnetoencephalographic (MEG) studies, a highly replicable finding is that during anticipatory attention, alpha-band (8-12 Hz) activity is modulated by attention-directing cues (Thut et al., 2006; Worden et al., 2000), and the patterns of alpha modulation encode the locus of spatial attention (Desantis et al., 2020; Foster et al., 2016; Lu et al., 2024; Rihs et al., 2007; Samaha et al., 2016; Trachel et al., 2015; Treder et al., 2011; van Gerven & Jensen, 2009). This is also true for slow potential shifts in event-related potentials (ERPs) and EEG/MEG signal patterns during anticipatory attention (Barnes et al., 2022; Harter et al., 1989; Hong et al., 2020; Hong et al., 2015; Hopf & Mangun, 2000; Thiery et al., 2016; Yamaguchi et al., 1994). In human fMRI studies, attention-directing cues result in increases in anticipatory baseline BOLD activity in thalamus (Woldorff et al., 2004) and retinotopic visual cortex (Giesbrecht et al., 2006; Hopfinger et al., 2000; Kastner et al., 1999; McMains et al., 2007; Park & Serences, 2022; Ress et al., 2000; Serences et al., 2004; Woldorff et al., 2004). Like in the animal studies of Luck and colleagues, the BOLD signals during anticipatory spatial attention are typically more robust in higher-order visual cortical areas (Hopfinger et al., 2000; O’Connor et al., 2002). In a more recent study, applying multivariate fMRI analysis techniques, Park and Serences (2022) found differences in multivoxel patterns evoked by different attention-directing cues in several retinotopic extrastriate visual areas, but as in prior studies, the effects during the anticipatory period were weaker in V1.

What is the role of baseline activity shifts in visual cortex during anticipatory visual attention? It is commonly thought that the stronger the top-down attentional modulation of the sensory cortex, the more effective would be the processing of the attended stimulus, which should then lead to better behavioral performance (Stokes et al., 2009). Ress et al. (2000) found that cue-related baseline BOLD increases in V1 through V3 were positively associated with task performance in a visual spatial attention task. Giesbrecht et al. (2006) reported similar findings for both spatial and feature attention. Fannon et al. (2008), however, found that cue-related pre-stimulus neural activity did not predict the stimulus-evoked activity or the subsequent behavioral performance. McMains et al. (2007) also reported that the post-stimulus effects of attentional selection observed in feature-selective regions were not accompanied by increased background activity during the anticipatory (pre-stimulus) period. Thus, there remains uncertainty regarding the physiological role and behavioral significance of the attentional modulations of baseline activity in visual cortex during anticipatory attention.

In order to shed new light on the physiological role and behavioral significance of baseline shifts in neural activity in visual cortex during anticipatory attention, we analyzed anticipatory period BOLD activity in a trial-by-trial spatial attention orienting task. We analyzed the signals across the entire visual cortical hierarchy from V1 through extrastriate cortex and posterior parietal cortex using both univariate and multivariate analysis methods. Further, we investigated the relationships between the strength of attentional biasing and behavioral performance. Although this study addresses a longstanding question in neuroscience, it provides advances over prior studies in two important respects. First, univariate, multivariate, and brain-behavior analyses were comprehensively carried out in the entire retinotopic visual hierarchy in a single study combining two datasets recorded at two different institutions using the same experimental paradigms. Second, in addition to the traditional cueing approach where different physical cues instructed the participants to covertly attend to different spatial locations, we included another condition in which the participants were allowed to freely choose which hemifield to attend when prompted by a choice cue or “prompt.” The inclusion of choice trials provides a strong test of the nature of top-down control over baseline activity in visual cortex that is free from the possible confounding influences of differences in the physical features of attention-directing cues.

## Methods

### Overview

Two datasets from studies conducted at the University of Florida (UF dataset) and the University of California, Davis (UCD dataset) were analyzed. By design, both datasets used the same experimental conditions. These datasets have been analyzed and published before to address different questions (Bengson et al., 2015b; Liu et al., 2016; Liu et al., 2017). None of the previously published studies examined how attention modulated baseline neural activity in retinotopic visual cortex.

*<u>UF Dataset</u>:* The Institutional Review Board at the University of Florida approved the experimental procedure. Eighteen healthy participants, between 18-22 years of age, provided written informed consent and took part in the experiment. Functional MRI data were collected from the participants while they performed a cued visual spatial attention task. Two participants were excluded from the analysis: one failed to follow the experimental instructions and another did not achieve behavioral accuracy above 70%. Three additional participants were removed due to excessive head/body movements. Data from the remaining thirteen participants was analyzed and reported here (n=13; 5 females and 8 males). EEG data were recorded along with the fMRI but not considered here.

*<u>UCD Dataset</u>:* The Institutional Review Board at the University of California, Davis, approved the experimental procedure. Nineteen healthy participants, between 18-22 years of age, provided written informed consent and took part in the experiment. Functional MRI data were recorded while the participants performed the visual spatial attention task. One participant was removed due to inconsistent behavioral performance (accuracy < 50% during the latter half of the experiment). Data from the remaining eighteen participants was analyzed and reported here (n=18; 2 females and 16 males).

We performed power analyses and found that: (1) For univariate analysis the required sample size to detect a significant effect (p<0.05 FDR) with a power of 0.8 was 22 (Durnez et al., 2016) and (2) for the MVPA decoding analysis the required sample size was 13. Thus, our study with a combined sample size of 31 is adequately powered to test our hypotheses. This sample size was also in line with the general recommendations for similar neuroimaging studies (Desmond & Glover, 2002; Pajula & Tohka, 2016). Similarly, our recent work applying both univariate and multivariate methods to fMRI data further suggests that there is sufficient power for carrying out the intended analyses (Bo et al., 2021; Meyyappan et al., 2021; Rajan et al., 2021). See below for our analytical strategy in which the two datasets were combined via meta-analysis.

### Experimental Paradigm

As shown in Figure 1, a fixation dot was placed at the center of the monitor and the participants were instructed to maintain eye fixation on the dot during the experiment. Two white dots in the lower left and lower right visual fields marked the two peripheral spatial locations where stimuli would be presented.

**Figure 1.**
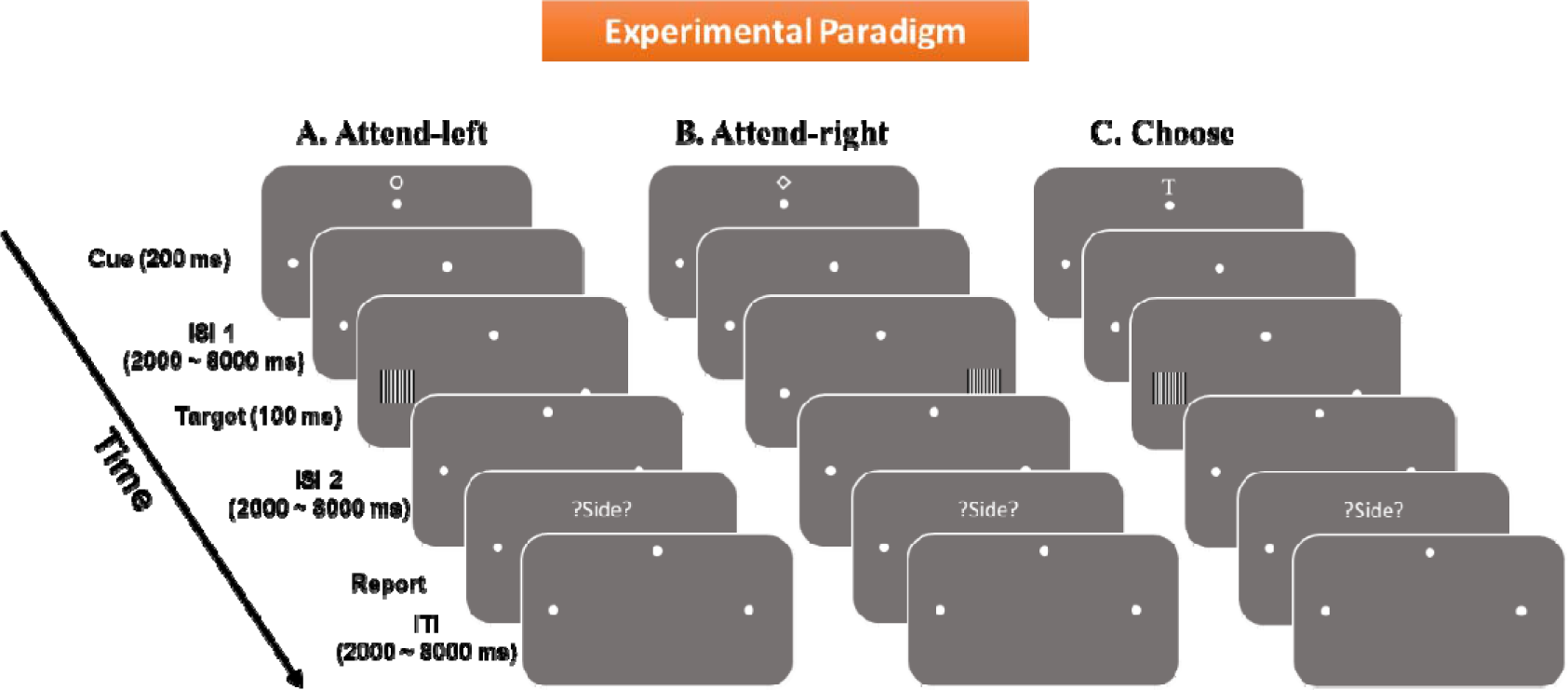
Experimental paradigm. Each trial started with one of three visual cues (200ms), two of them instructing subjects to covertly attend either left (A) or right (B) hemifields, indicated by the two white dots. A third cue, the choice cue (C), prompted the participants to choose either left or right visual field to attend. Following a variable cue-to-target delay (2000-8000ms), a target in the form of a grating patch appeared briefly for 100ms with equal probability in the left or the right hemifield. Participants were asked to discriminate the spatial frequency of the grating displayed in the cued or chosen location and ignore the un- cued or -un-chosen side. After a second variable ISI (2000-8000ms), participants were presented with a ‘?SIDE?’ screen and were asked to report the hemi-field they paid attention to in that trial. An inter-trial interval, varied randomly from 2000-8000ms, elapsed before the start of the next trial.

At the beginning of each trial, one of three symbolic cues (a letter T-shaped stimulus, a diamond, or a circle) was presented above the fixation dot for 200 ms. Two of the symbols explicitly instructed (cued) the participants to direct their attention covertly either to the left or right hemifield location (instructed trials) and the third prompted them to freely choose one of the two hemifields to covertly attend on that trial (choice trials). The three cue/prompt conditions occurred randomly and with equal probability, and the three symbols used as cues/prompts were counterbalanced as to their meanings across participants (i.e., cue-left, cue-right, and freely-choose). The choice condition, which was introduced originally to investigate willed attention (Bengson et al., 2015b; Liu et al., 2016; Liu et al., 2017; Rajan et al., 2019), was important to include for the purpose of this analysis in that it elicited the two attentional states (attend left vs. attend right) by using the same visual cue (the choice cue), thereby eliminating the influence of different visual cues on neural patterns in visual cortex (especially the early visual cortex), thus permitting the cross-validation of the findings from the instructional cue trials.

Following a random cue-target interval ranging from 2000 to 8000 ms, a black-and-white grating pattern (100% contrast) appeared at one of the two peripheral spatial locations for 100 ms and participants discriminated between two possible spatial frequencies of the target grating (0.2° per cycle vs. 0.18° per cycle for UF dataset; 0.53° per cycle vs. 0.59° per cycle for UCD dataset) appearing in the attended hemifield via a button-press and <u>ignored</u> the grating appearing in the unattended hemifield. The grating occurred in either the left or right visual hemifields with <u>equal probability</u>, in other words, there was a 50% chance that the stimulus appeared at the attended location, while in the remaining 50% of the trials, the stimulus appeared at the un-attended location. According to this design, withholding response during un-attended trials and accurately identifying the spatial frequency of the stimulus during attended trials were both deemed correct responses. After the target stimulus, following a variable inter-stimulus interval (ISI) ranging from 2000 to 8000 ms, participants were prompted by the visual cue “?SIDE?” to report the spatial location that they attended on that trial. This reporting was needed to track the participant’s direction of attention in the choice attention condition and was also included in instructed trials to maintain consistency across conditions. A variable inter-trial interval (ITI; 2000-8000 ms) was inserted after the subject reported the side attended.

Reporting of the side attended was highly accurate across the group of participants, with the accuracy being 98.4%, averaged across conditions. While side-reporting accuracy could be determined by comparing self-reports with the cue during instructed attention trials, the accuracy for choice attention trials was further determined by examining the consistency between participants’ responses to targets and the self-reports (e.g., responded to the target in the left visual field and reported choosing left).

For the choice or willed attention condition, participants were directed to make impromptu (spontaneous) decisions regarding which side to attend when presented with a choice cue and advised against adhering to any stereotypical strategies such as consistently attending to the same or opposite side as in the previous trial. Choose-left (UF: 50.57% ± 3.17%; UCD: 47.88% ± 1.84%) and choose-right (UF: 49.43% ± 2.69%; UCD: 52.12% ± 1.84%) trials were evenly distributed and not significantly different from each other (UF: p = 0.86; UCD: p = 0.26).

All participants went through a training session prior to scanning to familiarize them with the task and to ensure adequate performance (above 70% correct) and compliance with the procedures (e.g., maintain fixation to the center throughout the experiment). Though no eye-tracking was performed in the scanner at UF, at UC Davis the study team monitored participants for eye movements using an MR-compatible Model 504 Applied Sciences Laboratories eye-tracker. The experiment was divided into blocks of trials with the duration of each block being approximately six minutes (60 trials).

### Data Acquisition

*UF Dataset:* MR images were acquired at the University of Florida using a 3T Philips Achieve scanner equipped with a 32-channel head coil. Functional magnetic resonance images (fMRI) were obtained using an echo-planar sequence (EPI) with parameters as follows: repetition time (TR) = 1980ms; echo time (TE) = 30ms; matrix = 64×64 field of view = 224mm, slice thickness = 3.5mm. Thirty-six transverse slices parallel to the plane connecting anterior to posterior commissure were obtained in ascending order with a voxel resolution of 3.5×3.5×3.5mm.

*UCD Dataset:* MR images were collected at the Imaging Research Centre of the University of California at Davis using a 3-T Siemens Skyra scanner with a 32-channel head coil. Functional magnetic resonance images were obtained using an echo-planar sequence with parameters as follows: TR = 2100ms; TE = 29ms, matrix = 64×64, field of view = 216mm, slice thickness = 3.4mm. Thirty-four transverse slices were obtained within an interleaved acquisition order and slices were oriented parallel to the plane connecting the anterior-posterior commissure. The voxel resolution was 3.5×3.5×3.5mm.

The slight differences in scanning parameters were due to optimization done at each recording site. In particular, the simultaneous recording of EEG and fMRI at UF necessitated the setting of parameters that were more conducive to the removal of scanner artifacts from the EEG data.

### Data Preprocessing

The fMRI data were preprocessed in SPM with custom-created MATLAB scripts. The slice timing differences were corrected with a *sinc* interpolation to account for the acquisition delay. Motion realignment was performed by co-registering all images with the first scan of each session. Six motion parameters (3 translational and 3 rotational) were calculated and regressed from the fMRI data. All images were spatially co-registered to the standard MNI space and resampled to a voxel resolution of 3×3×3mm. Finally, the spatially normalized images were smoothed using a Gaussian kernel of 8mm full width at half maximum. A high-pass filter with a cutoff frequency of 1/128Hz was used to remove any low-frequency noise, and global signals were removed by adjusting the intensities of each voxel(Fox et al., 2009).

### Whole-brain Univariate Analysis

Voxel-level univariate BOLD activation analysis was carried out using the general linear model (GLM) method where an experimental event (e.g., cue onset) is assumed to evoke a stereotyped hemodynamic response (i.e., the hemodynamic response function or HRF). Seven regressors were used to model the experimental events, including two for instructed attention conditions (attend-left and attend-right), two for choice attention conditions (choose-left and choose-right), one for incorrect trial, two for stimuli appearing in the left and right visual field. The cue-evoked activation was obtained within each subject by contrasting the appropriate regressors using a t-test for each voxel. The group-level activation map was obtained by performing a one-sample t-test on each subject’s contrast maps with a threshold of p<0.05 correcting for multiple comparisons with False Discovery Rate (Nichols & Hayasaka, 2003).

### Estimation of Single-trial BOLD Activity

The MVPA approach is applied at the single trial level. All trials with correct responses were used. A trial was considered correct if either no response was made during the unattended trials or if the spatial frequency of the stimulus was accurately identified during the attended trials. To estimate the trial-by-trial cue-evoked BOLD activity we applied the beta series regression method (Rissman et al., 2004). Specifically, cue-evoked BOLD activity was modeled using a separate regressor for each instructed and choice trial in the general linear model (GLM), and one additional regressor was used for all the targets to account for the target-evoked activities and to remove their possible influences on the estimation of cue-evoked activity. The coefficient of each regressor, referred to as the beta value, reflected the cue-related BOLD response for that trial.

### Region of Interest (ROI) Selection

Sixteen retinotopic visual regions from a probabilistic visual topography map by Wang et al. (2015) were used as regions of interests (ROIs). Ordered from low to high in the visual hierarchy (dorsal and ventral pathways combined), they are: V1v, V1d, V2v, V2d, V3v, V3d, hv4, V3a, V3b, LO1, LO2, VO1, VO2, IPS, PHC1, and PHC2.

### Multivoxel Pattern Analysis (MVPA)

The MVPA method uses pattern classification algorithms to distinguish between two brain states based on distributed BOLD activities across multiple voxels in a given ROI. In this study, we used support vector machine (SVM) in the MATLAB Statistics and Machine Learning Toolbox as the classifier for the MVPA analysis. A 10-fold cross-validation approach was applied to minimize the effects of overfitting. The trial assignment into each fold was fully random. The classifier performance was measured in terms of decoding accuracy. Above-chance decoding accuracy is taken to signify the presence of attention-related signals in the ROI, and the higher the decoding accuracy, the larger the difference between the two BOLD activation patterns evoked by the two attention conditions (attend left vs attend right), which is hypothesized to indicate stronger top-down attention modulation. The cross-validation analysis was repeated 25 times over different random fold partitions to avoid any bias that may have resulted when grouping the trials into 10 specific groups (10-fold). Twenty-five,10-fold cross-validation accuracies (a total of 250 decoding accuracies) were averaged to obtain the decoding accuracy for a subject.

For statistical analysis, the decoding accuracy was compared against the chance level accuracy of 50% using one-sampled t-tests at p<0.05, corrected for multiple comparisons using FDR.

### Relating Decoding Accuracy with Behavior

To understand the functional significance of top-down biasing of visual cortex, the decoding accuracy at the subject level and behavioral performance at the subject level in target discrimination was correlated across subjects. Given that there are 16 ROIs representing all the areas within the retinotopic visual hierarchy, and expecting that these decoding accuracies was likely to covary to a large degree, we performed a principal component analysis (PCA) on the decoding accuracies from the 16 ROIs for UF and UCD datasets to extract the underlying common variance and correlated (Spearman’s rank correlation) the score on the first PCA component with behavior. Then, for completeness, each ROI’s relation between decoding accuracy and behavior was further tested.

The behavioral performance was characterized by the efficiency score (ES), which is the ratio of accuracy over reaction time (accuracy/RT), with a higher score indicating better behavioral performance. Prior work has used the inverse efficiency score (Bruyer & Brysbaert, 2011), which is the ratio of reaction time over accuracy, to measure behavioral performance, with a lower score indicating better behavioral performance. We note that in our design, the instruction to the participants was to respond *as fast as they could* while also being *as accurate as they could*, the speed-accuracy tradeoff may prevent the use of RT or accuracy singly as the appropriate behavioral measure. Computing the ratio of accuracy and RT can account for the speed-accuracy trade-off effect while retaining the main task effect (Bruyer & Brysbaert, 2011; Liesefeld & Janczyk, 2019; Vandierendonck, 2017).

### Combining Datasets Using Meta-analysis

Analysis was done first on the UF and UCD datasets separately, we then combined the two datasets through meta-analysis to enhance statistical power. An effect was considered statistically significant if it is significant in the meta-analysis. The meta-analysis was done using the Lipták-Stouffer method (Lipták, 1958). This method has been used commonly to combine results from different datasets (Cheng et al., 2015; Huang & Ding, 2016). It involves converting the p-value of each dataset to its corresponding Z value using the formula:

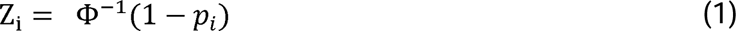

Here Φ denotes the standard normal distribution function. The Z values of different datasets were then combined using the Liptak-Stouffer formula:

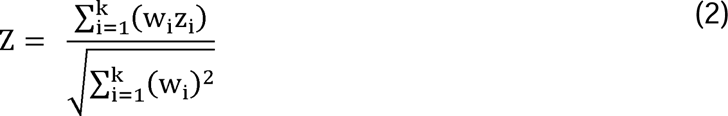

Here Z is the combined z score of k datasets (for the present study k=2), i refers to the i^th^ dataset or site, w_i_ is the weight i^th^ dataset which is equal to √*N_i_*, where *N_i_* is the number of samples in i^th^ dataset. From the combined Z score, the corresponding p-value can then be assessed. Combining data from multiple datasets using meta-analysis improves the statistical power by reducing the standard error of the weighted average effect-size. A smaller standard error results in a narrower confidence interval around the estimated effect, increasing the likelihood of detecting true effects (Cohn & Becker, 2003).

## Results

### Behavioral Analysis

***UF dataset:*** For instructed trials, target discrimination accuracy for cue-left trials and cue-right trials was 85.72 ± 2.1% and 85.80 ± 1.2%, respectively (Figure 2A); there was no statistically significant difference between the two conditions (t(12) = −0.03, p = 0.96, d = −0.01). Reaction time for cue-left trials was 918.56 ± 24.49, and for cue-right trials was 911.81 ± 23.28 ms (Figure 2B); there was no statistically significant difference between the two conditions (t(12) = 0.43, p = 0.67, d = 0.07). For choice trials, accuracy was 88.41% ± 1.9% for choose-left and 90.78% ± 1.3 % for choose-right trials (Figure 2A), and the reaction time was 942.27 ± 47.75 ms for choose-left and 938.25 ± 29.04 ms for choose-right trials (Figure 2B). There was no statistically significant difference between choose-left and choose-right for either accuracy or reaction time (t_acc_ (12) = −1.2, p_acc_ = 0.25,d_acc_= −0.39; t_RT_ (12) = 0.09, p_RT_ = 0.92, d_RT_ = 0.03). A one-way main effects ANOVA further confirmed that there were no significant differences in behavioral performance between instructed and choice trials [F_acc_ (3,12) = 2.07, p_acc_ = 0.11, η_acc_^2^ = 0.1; F_RT_ (3,12) = 0.21, p_RT_ = 0.89, η_RT_^2^ = 0.01; Figure 2A and 2B].

**Figure 2.**
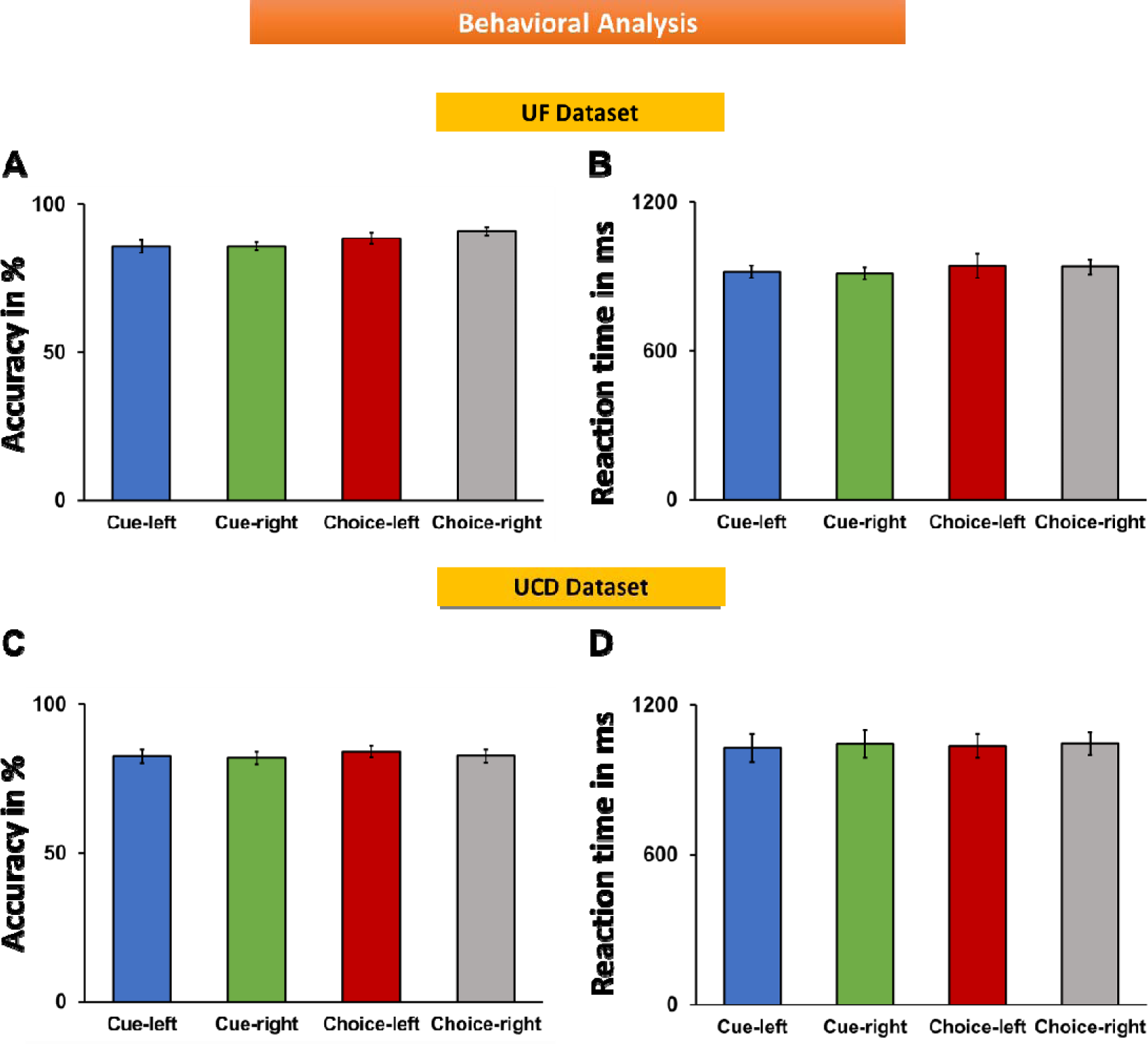
Behavioral results from instructed and choice trials. (A & C): Comparison of accuracy (ratio of correct trials and total number of valid trial’s) between different attention conditions for UF (A) and UCD (C) datasets. (B & D): Comparison of reaction times (RT) between different attention conditions for UF dataset (B) and UCD dataset (D).

***UCD dataset:*** For instructed trials, response accuracy for cue-left trials and cue-right trials was 82.45 ± 2.3 % and 81.97 ± 2.1% (Figure 2C); there was no statistically significant difference between the two conditions (t(17) = 0.43, p = 0.67, d = 0.05). Reaction time for cue-left trials was 1027.02 ± 55.60 ms and for cue-right trials 1041.87 ± 54.95 ms (Figure 2D); there was no statistically significant difference between the two conditions (t(17) = −0.68, p = 0.50, d = −.06). For choice trials, the accuracy was 84.07% ± 2.0% for choose-left and 82.60% ± 2.1% for choose-right trials, and the reaction time was 1034.42 ± 48.06 ms for choose-left and 1043.43 ± 45.84 ms for choose-right trials. There was no statistically significant difference between choose-left and choose-right in either accuracy or reaction time (t_acc_(17) = 0.98, p_acc_ = 0.34, d_acc_ = 0.16; t_RT_(17) = −0.36, p_RT_ = 0.72, d_RT_ = −0.04). A one-way ANOVA further confirmed that there were no significant differences in behavioral performance between instructed and choice trials [F_acc_ (3,17) = 0.2, p_acc_ = 0.90, η_acc_^2^ = 0.008; F_RT_ (3,17) = 0.02, p_RT_ = 0.99, η_RT_^2^ = 0.01; Figure 2C and 2D].

### Univariate Analysis of Cue-Evoked BOLD Activation

Two types of univariate analyses were carried out. First, general linear model (GLM) analysis was performed at the whole brain level. For instructed trials, the attention-directing cues activated the dorsal attention network (compared to baseline), including frontal eye field (FEF) and intraparietal sulcus/superior parietal lobule (IPS/SPL), as well as higher-order visual areas such as superior occipital cortex, for both UF and UCD datasets (Figure 3A-3D), in line with previous studies (Giesbrecht et al., 2003; Heinze et al., 1994; Hopfinger et al., 2000). Contrasting attention-directing cues (cue-left vs. cue-right) revealed no activations at p<0.05 (FDR) (not shown). Some higher-order visual areas, including lingual and fusiform cortex, began to appear at a much-relaxed threshold of p<0.01 (uncorrected) (Figure 3E-3H), suggesting that attended information is not strongly encoded in ROI-level univariate signals. Similar results were also found for choice trials (not shown).

**Figure 3.**
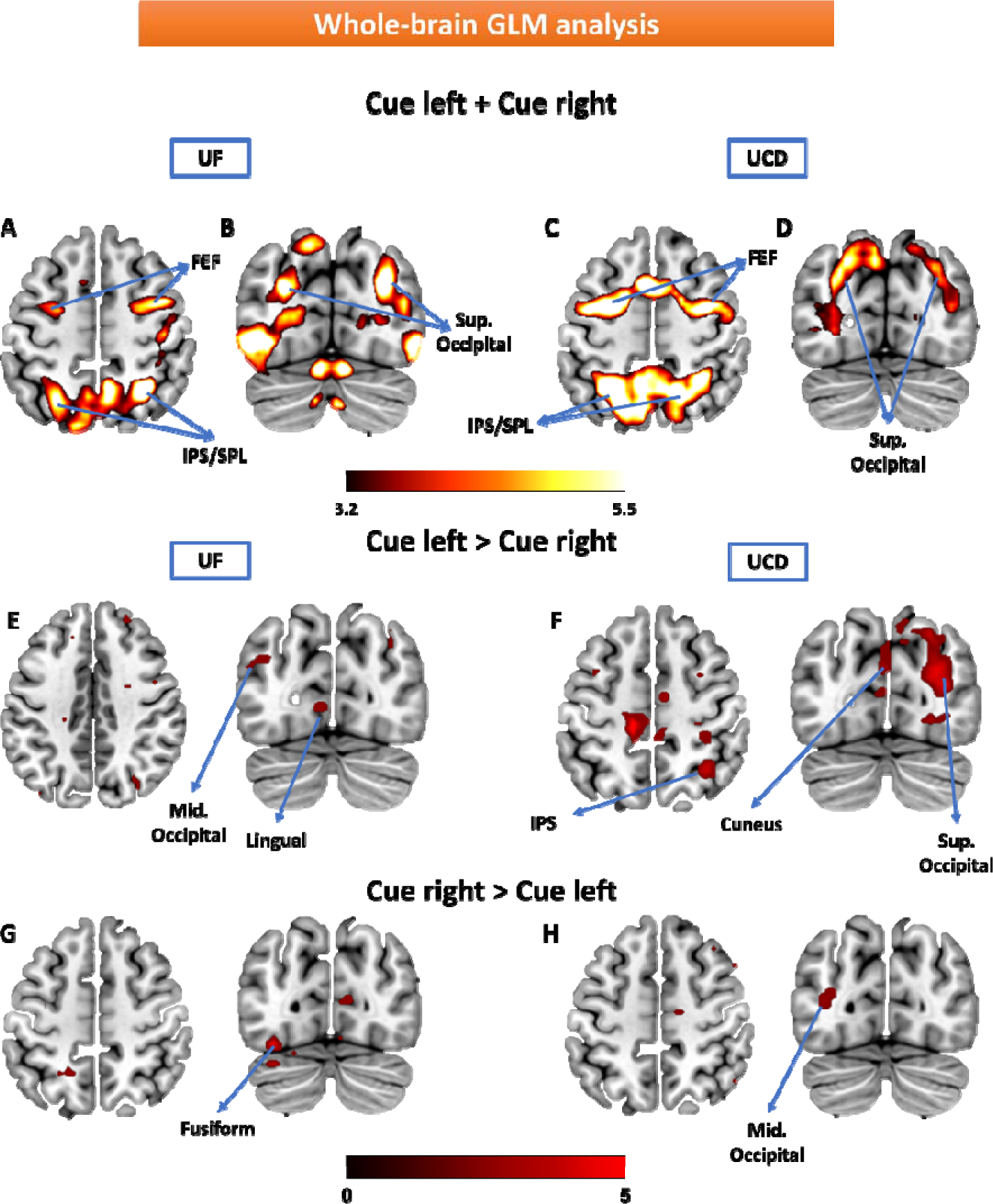
Whole-brain GLM analysis of cue-evoked BOLD activity. Attention cues (left and right combined) evoked significant activation in dorsal attention network (FEF,IPS/SPL) and extra-striatal visual cortex (p<0.05 FDR corrected) for both UF (A & B) and UCD (C & D) datasets. However, contrasting attention-directing cues (cue-left vs cue right) revealed no activation when thresholded at p<0.05 with FDR correction (not shown). High-order visual regions only began to appear after lowering the threshold to p<0.01 uncorrected: cue-left > cue-right (E & F) and cue-right > cue-left (G & H).

Next, to investigate the univariate activity at the ROI level in the retinotopic visual cortex, the BOLD data in the hemisphere when it was ipsilateral to the cued direction was subtracted from that in the same hemisphere when it was contralateral to the cued direction (i.e., cue-left minus cue-right for right hemisphere, and cue-right minus cue-left for left hemisphere) and the difference was averaged across the two hemispheres and within each ROI. As shown in Figures 4A and 4B, for higher-order visual cortex such as LO and IPS, covert attention elicited significant univariate BOLD activation, and the effects were consistent across the two datasets, in line with previous studies (Giesbrecht et al., 2003; Heinze et al., 1994; Hopfinger et al., 2000). For early visual areas such as V1 and V2, however, the effects were less consistent across the two datasets. A meta-analysis combining the two datasets showed that there were no significant attention-specific <u>univariate</u> neural modulations in early visual areas except V1v (Figure 4C). To test whether ROI-level univariate activation exhibited systematically increasing tendency as we ascended the visual hierarchy, a linear regression analysis was applied, in which univariate activation for each ROI was treated as a function of the ROIs, and the ROIs were given numbers ranging from 1 to 16. A marginally significant positive slope in the meta-analysis (p = 0.07) suggested the possibility of such a tendency. When correlating the univariate activation difference for each ROI with behavioral efficiency (response accuracy/reaction time), combining the two datasets via meta-analysis, none of the regions in the visual hierarchy exhibited a significant correlation between attention-related univariate BOLD activity and behavior (p>0.05 FDR; Figure 4D).

**Figure 4.**
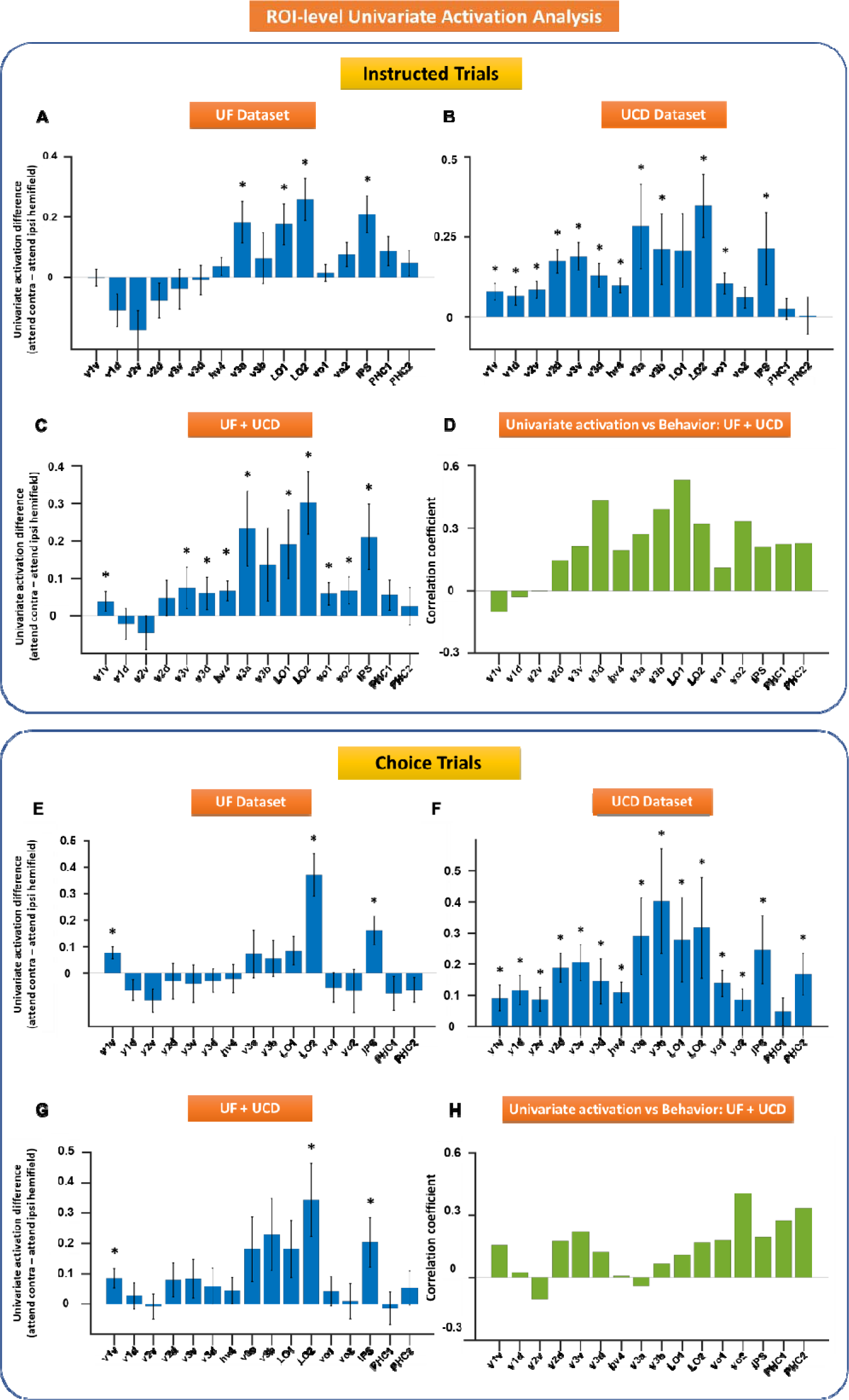
ROI-level analysis of cue-evoked BOLD activity. BOLD activity difference between attention to contralateral and ipsilateral visual fields were computed and averaged across the hemispheres for instructional (A-D) and choice trials (E-H) for both datasets: UF (A & E) and UCD (B & F). (C & G) activation differences from UF and UCD datasets were averaged from instructed (C) and choice trials (G) and the respective p values obtained via meta-analysis. (D & H) correlation between behavior and activation differences across ROIs from instructed (D) and choice trials (H). Here the correlation coefficients are averaged across the datasets and the p values from both the datasets were combined using meta-analysis. The error bars denote the standard error of the mean (SEM). * p < 0.05 FDR.

Applying the same analysis to the choice trials, we found similar results (Figure 4E-4G). As with instructed trials, for choice trials, attention modulations of univariate BOLD activity were mainly found in higher order visual areas with V1v being the notable exception. While Figures 4C and 4G exhibited similar patterns, a linear regression analysis combining the two datasets did not yield a statistically positive slope (p = 0.13). The smaller number of choice trials is likely a factor weakening the statistical power. Further, no significant correlations were found between univariate attention-related BOLD activation and behavioral performance (Figure 4H).

### Multivariate Analysis of Cue-Evoked BOLD Activation: Instructed Trials

Multivoxel pattern analysis (MVPA) was applied to each ROI in each of the two datasets. The decoding accuracy between cue-left and cue-right was above the chance level in all visual regions in both UF (Figure 5A) and UCD datasets (Figure 5B). Meta-analysis combining the two datasets confirmed these findings, suggesting that the top-down biasing signals are present in all areas of the visual hierarchy. In addition, the decoding accuracy tended to be higher in lower-order visual areas than in higher-order areas (Figures 5C & 5D), in contrast to the pattern of univariate activation seen in Figure 4. A linear regression analysis treating decoding accuracy for each ROI as a function of the ROIs yielded a significantly <u>negative</u> slope for both datasets with the meta-analysis p-value being p = 4.7 x 10^−7^. To further demonstrate the consistency of the results between the two datasets, for each ROI, we plotted the decoding accuracy averaged over all participants from the UF dataset against that from the UCD dataset in Figure 5E. The Spearman rank-correlation across all 16 ROIs was highly significant with r = 0.75 (p = 0.001), suggesting a high degree of similarity in the decoding accuracy distribution across visual cortical regions between the two datasets.

**Figure 5.**
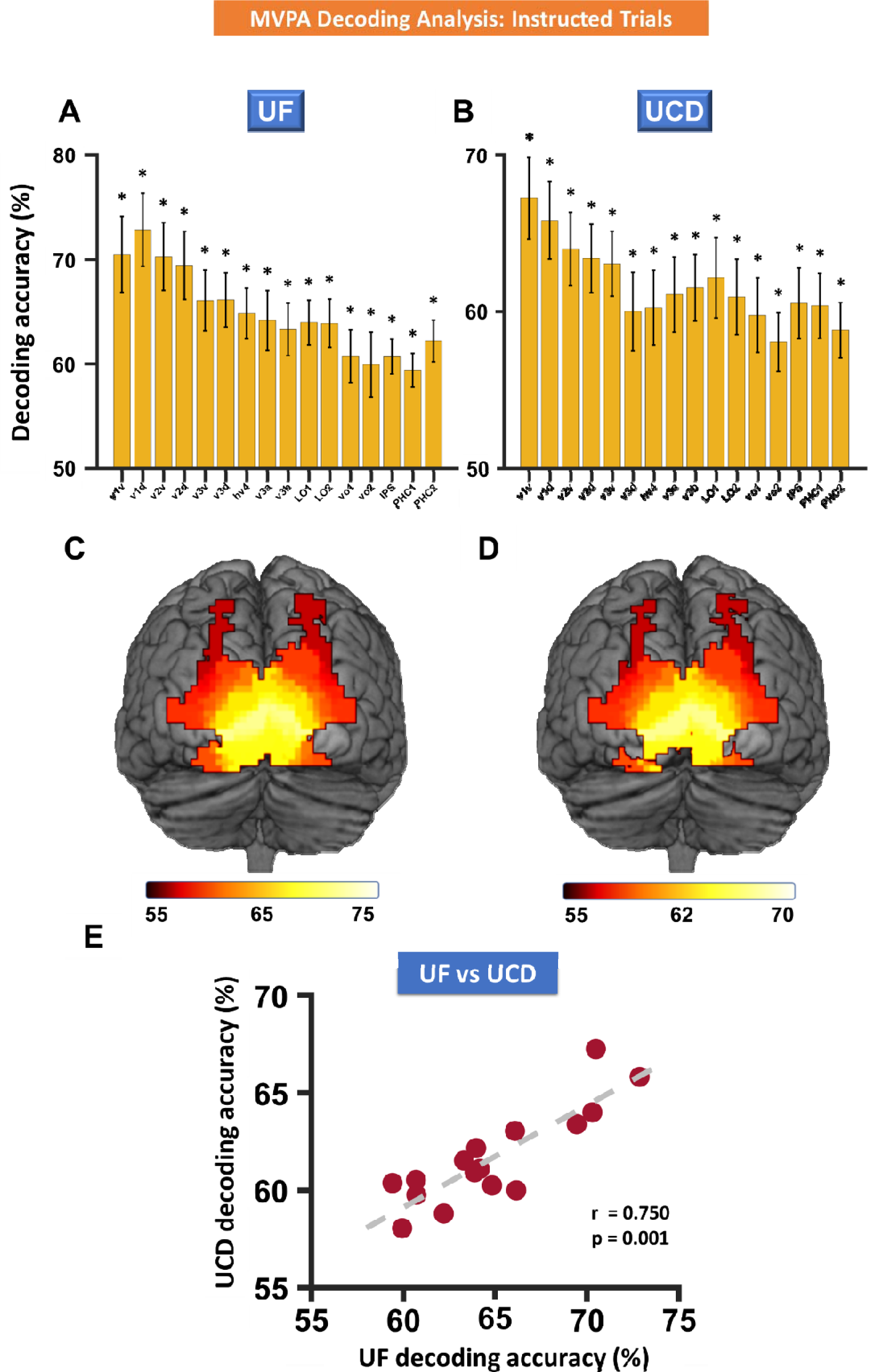
Multivariate pattern analysis of cue-left vs cue-right: Instructed trials. (A & B) Decoding accuracies in different visual ROIs for UF dataset (A) and UCD dataset (B). (C & D) Posterior view of color-coded decoding accuracies for UF dataset (C) and UCD dataset (D). (E) Scatter plot comparing UCD vs. UF decoding accuracies where each dot represents a visual ROI. The error bars denote the SEM. * p < 0.05 FDR

The decoding accuracy reflects the degree of distinctness of the neural activity in the two attention conditions (cue-left vs cue-right). The higher the decoding accuracy, the larger the difference between the two neural patterns underlying the two attention conditions, which was taken to indicate stronger attention modulation. We thus used decoding accuracy as a quantitative index of the strength of top-down biasing to evaluate the relationship between top-down attention control and behavioral performance. Because decoding accuracy across different ROIs exhibited a high degree of correlation across participants (not shown), indicating that the variance in the decoding accuracy across ROIs can be captured by fewer latent variables, a principal component analysis (PCA) was performed to reduce the dimensionality of 16 ROI-based decoding accuracies. In particular, the first PCA component explained 79.75% and 83.49% of the variance for UF and UCD datasets respectively (Figure 6A and 6B). Thus, each subject’s score on the first PCA component was used to represent the magnitude of decoding accuracy for that subject and correlated with the z-scored behavioral efficiency (response accuracy/reaction time). A significant positive correlation between decoding accuracy and behavioral performance was found for both datasets; UF Dataset: r = 0.5824, p = 0.0403 (Figure 6C) and UCD Dataset: r = 0.6780, p = 0.0026 (Figure 6D). Combining the two datasets using meta-analysis showed that the overall correlation effect was robust and highly significant (p = 0.0005). At the ROI level, decoding accuracy is significantly correlated with behavior across all the retinotopic regions within the visual hierarchy; see Figure 6E for ROI-level correlation coefficients averaged across the two datasets (meta-analysis p < 0.05 FDR). There are no clear differences in the brain-behavior relationship between lower-order and higher-order visual areas. These findings suggest that the stronger the top-down biasing signals in the retinotopic visual regions, indexed by higher decoding accuracy, the better the behavioral performance (Stokes et al., 2009).

**Figure 6.**
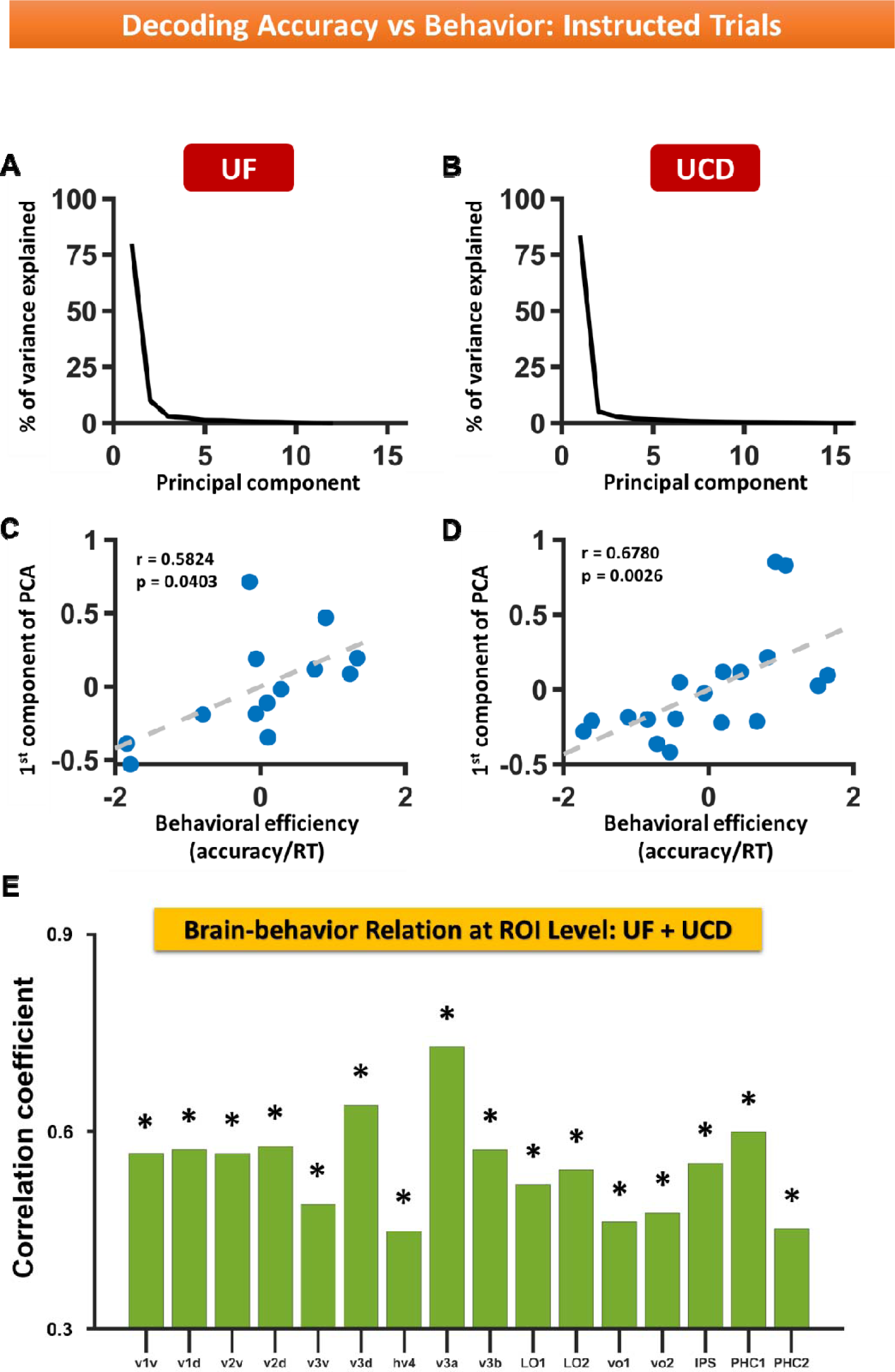
Decoding accuracy vs behavior: Instructed trials. (A & B) Percentage of variance explained by the principal components of decoding accuracies across ROIs for (A) UF and (B) UCD datasets. (C & D) Decoding accuracy represented by score on first PCA component vs behavioral efficiency (z-score) for (C) UF and (D) UCD datasets. (E) Correlation between decoding accuracy and behavior across ROIs. Here the correlation coefficients are averaged across the two datasets and the p values from both the datasets were combined using meta-analysis; the correlation is significant for all ROIs at p < 0.05 FDR.

### Multivariate Analysis of Cue-Evoked BOLD Activation: Choice Trials

In instructed trials, the two attentional states (attend-left vs attend-right) were the result of two different symbolic instruction cues. The decoding analysis in the visual ROIs especially in the early visual ROIs could be influenced by the physical differences in the two symbolic cues. The inclusion of choice trials where the two attentional states resulted from the same cue (choice cue) helped to address this problem. As shown in Figure 7A, where the two datasets were combined, decoding choose-left versus choose-right yielded decoding accuracies that were significantly above chance level in all ROIs (meta-analysis p<0.015 FDR). It is worth noting that because we had half as many choice trials as instructed trials (cue-left, cue-right and freely choose cues each accounted for 1/3 of trials), there was less statistical power to detect differences between attention conditions for choice trials. Figure 7B shows that the decoding accuracy distribution across the ROIs was consistent across the two datasets (cf. Figure 5E). Across the visual hierarchy, similar to the instructed trials, the linear regression analysis yielded a significantly <u>negative</u> slope combining the two datasets (meta-analysis p = 0.023), showing that the decoding accuracy decreases as we ascend the visual hierarchy.

**Figure 7.**
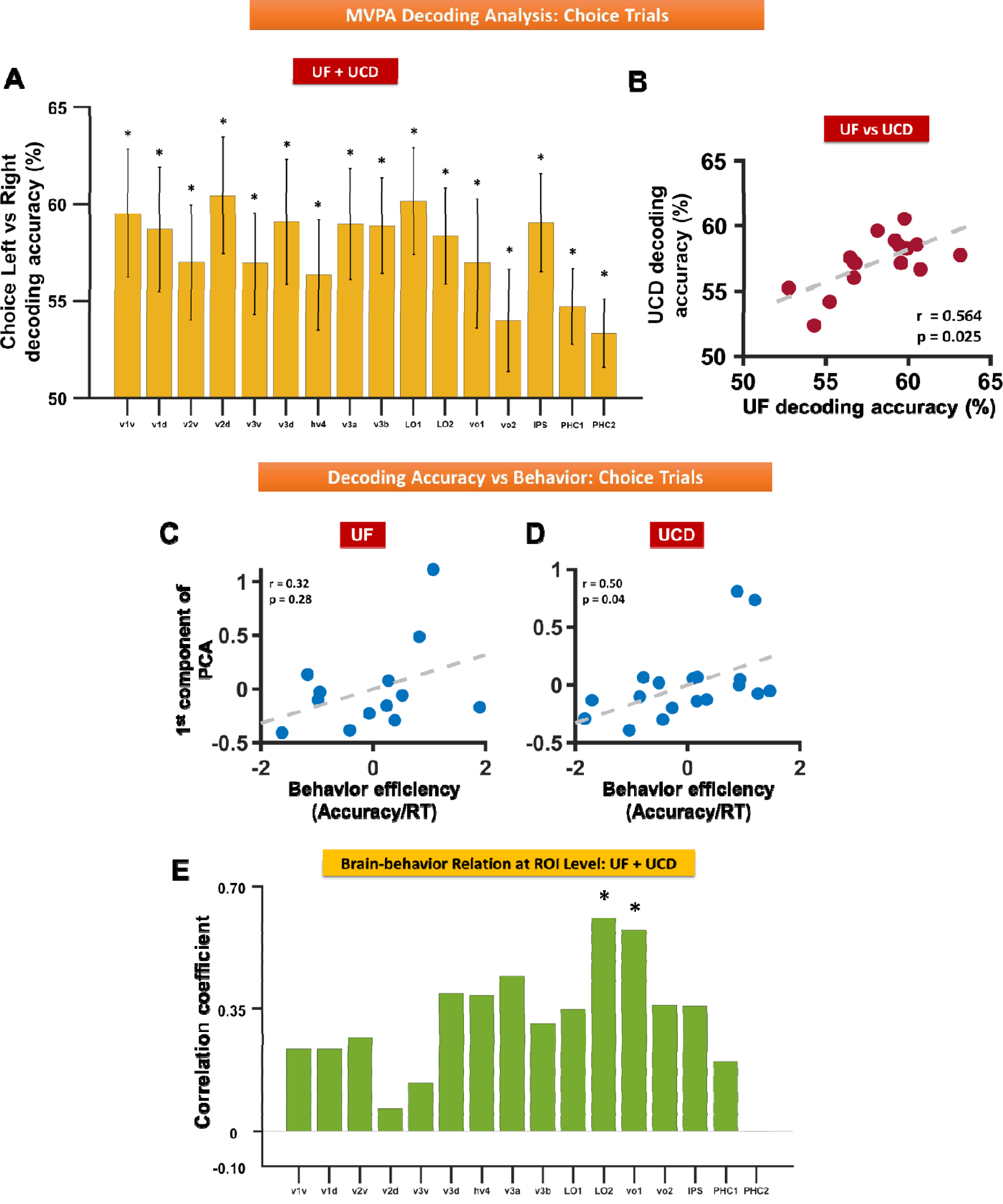
Decoding accuracy for attend-left vs attend-right for choice trials. (A) Decoding accuracies averaged from UF and UCD datasets for different visual ROIs; p values were determined via meta-analysis. (B)Scatter plot comparing UCD vs. UF decoding accuracies where each dot represents a visual ROI. (C & D) Decoding accuracy vs. behavioral efficiency for UF dataset (C) and for UCD dataset (D). (E) Correlation between decoding accuracy and behavior across visual-topic ROIs. The error bars denote the SEM. * p < 0.05 FDR.

Next, we tested whether the decoding accuracies predict behavioral performance. The first PCA component of the 16 decoding accuracies (16 ROIs) explained 73% and 66% of the variance for the UF and UCD datasets respectively. Correlating the score on the first PCA component with behavioral efficiency (response accuracy/reaction time), we found a significantly positive correlation for the UCD dataset: r = 0.5004, p = 0.0363 (Figure 7D) and a positive but not statistically significant correlation for the UF dataset: r = 0.3242, p = 0.2799 (Figure 7C). When the two datasets were combined, the p-value from the meta-analysis was p = 0.0361. suggesting a significant positive correlation. This decoding accuracy-behavior result from the choice trials is in agreement with the decoding accuracy-behavior result from the instructed trials; namely, the more distinct the BOLD patterns underlying the two attention conditions in the retinotopic regions, the stronger the top-down biasing signals in these regions, and the better the behavioral performance. Across visual ROIs, the decoding accuracy-behavior correlation is significant in higher-order areas VO1 and LO2 (Figure 7E). For the other ROIs, while a positive correlation was seen, the correlation values were not able to pass the statistical threshold of p<0.05 (FDR). Here, as indicated earlier, the smaller number of choice trials is again a factor that has contributed to weaken the statistical power to detect the brain-behavior relation in some of the ROIs.

### Comparing Instructed and Choice Trials

The neural patterns underlying the two attentional states (attend-left vs attend-right), whether the result of instruction cues or the choice, were expected to share commonalities. Three analyses were carried out combining the two datasets to compare instructed and choice trials. First, the instructed and choice decoding accuracies (averaged across the two datasets) were compared in Figure 8A, and a significant correlation was found (r = 0.57 and p = 0.02), suggesting that each ROI was similarly modulated by top-down attention whether the attentional states were formed from instructional cues or by pure internal decisions of the subjects. Second, for each dataset, we utilized the method of cross-decoding, by taking the SVM classifier trained on **instructed** trials and testing it on **choice** trials. We found the cross-decoding accuracies averaged across the two datasets (Figure 8B) were significantly above chance level in all the ROIs (meta-analysis p<0.05 FDR), suggesting that the patterns of neural activities resulting from instructional cues and from internal choices reflected attention modulations and not significantly influenced by the physical properties of the cues. Third, cross-decoding accuracy (model trained on instructed trials and tested on choice trials) and self-decoding accuracy (model trained on choice trials and tested on choice trials) were averaged across the two datasets and compared and the differences were not statistically significant (meta-analysis p > 0.05 FDR; Figure 8C), further confirming that the pattern of neural activity observed in the retinotopic visual cortex reflects top-down attentional biasing signals, and not merely the differences in the stimulus properties of the two cues in instructed trials.

**Figure 8.**
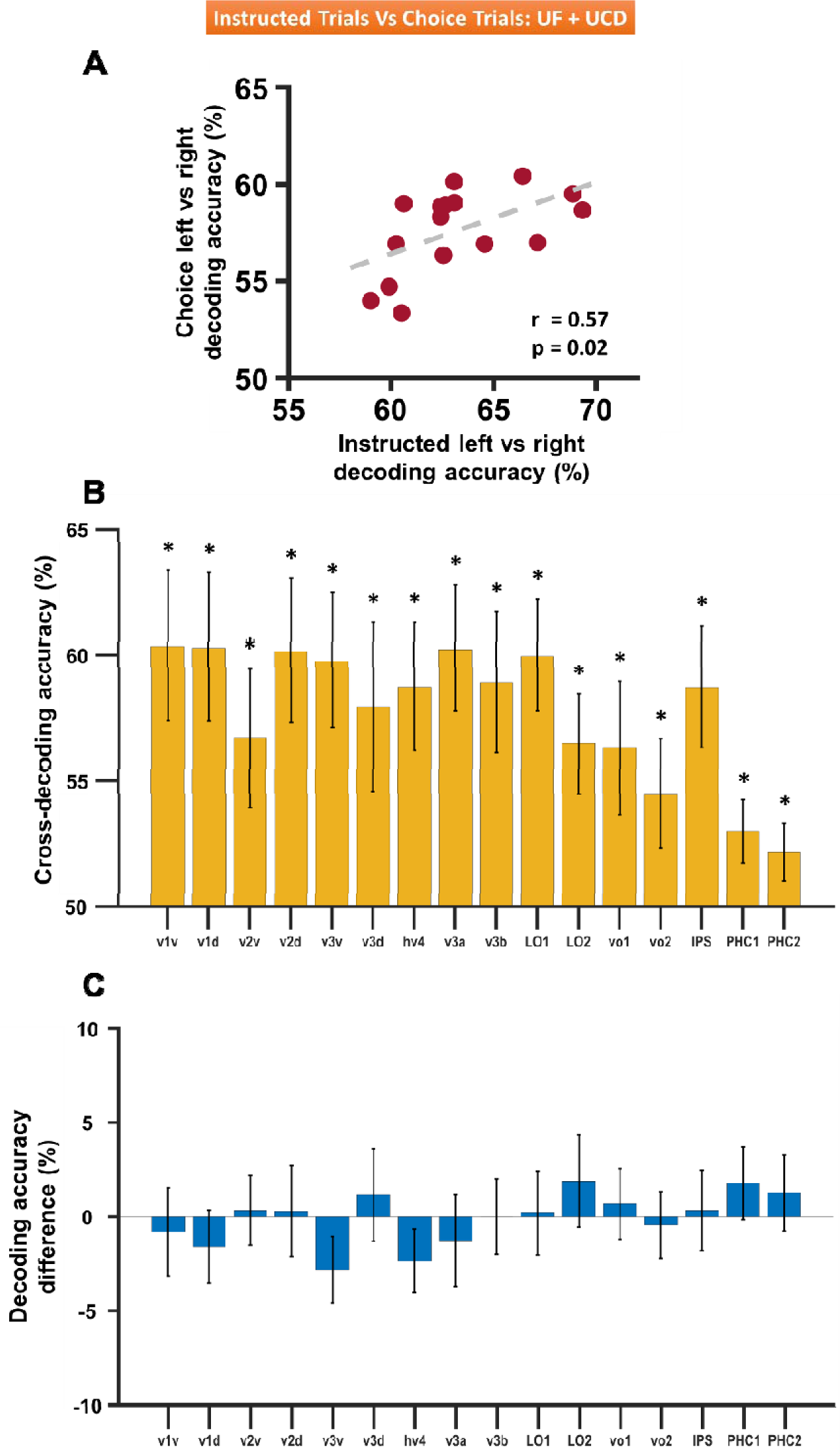
Comparison between choice trials and instructed trials (UF and UCD datasets combined). (A) Choice-left vs choice-right decoding accuracies plotted against cue-left vs cue-right decoding accuracies; each dot represents a visual ROI. (B) Cross decoding accuracy comparison across ROIs where classifiers were trained on instructed trials and tested on choice trials. (C) Difference between cross-decoding accuracy (classifiers trained on instructed trials and tested on on choice trials) and self-decoding accuracy (classifiers trained on choice trials and tested on choice trials) for different ROIs. The error bars denote the SEM. * p < 0.05 FDR.

## Discussion

We considered two questions in the control of visual spatial attention. First, what is the distribution of top-down biasing signals across different identified visual areas within the visual cortical hierarchy during spatial attention? More specifically, are early visual areas such as V1 more weakly biased during anticipatory spatial attention as has often been reported (e.g., O’Connor et al., 2002)? Second, what is the functional role of top-down control signals? Most models hold that they result in neuronal biasing that facilitates subsequent selective target processing (e.g., Chawla et al., 1999). If so, is the magnitude of the observed biasing signals during anticipatory attention predictive of behavioral performance? The answer to this question has remained unclear in the literature(Fannon et al., 2007; Zheng et al., 2024).

To answer these questions, we analyzed fMRI data from two different studies in which subjects performed a trial-by-trial visual spatial attention task. Two different methods of triggering the orienting of spatial attention were included: one involved the use of instructional cues to direct the subject’s attention to the left or the right visual field, and the other involved the use of a choice cue that prompted the subject to make voluntary decisions on a trial-by-trial basis about whether to attend the left (choose-left) or the attend the right (choose-right) visual field(Bengson et al., 2015a; Bengson et al., 2014). Applying both univariate and multivariate techniques to analyze cue-related neural activity (i.e., baseline shifts) in the retinotopic visual cortex, we obtained the following results.

Among sixteen retinotopic visual ROIs, differential attend versus ignore <u>univariate</u> BOLD activations were found in higher-order visual areas but only weakly or not at all in lower-order visual areas (i.e., V1v being the exception). These differential <u>univariate</u> activations in visual cortex during the anticipatory period failed to predict task performance. In sharp contrast, <u>multivoxel patterns</u> differentiated the two attention states (attend left vs. attend right) in all 16 ROIs, with lower-order ROIs (e.g., V1) having <u>higher</u> decoding accuracy than higher-order visual ROIs. The more distinct the attend-left versus attend-right multivariate neural patterns (i.e., the higher the decoding accuracy) in the visual cortex, the better the subject’s behavioral performance measured by behavioral efficiency - accuracy/reaction time(Liesefeld & Janczyk, 2019; Townsend & Ashby, 1983). The neural patterns associated with instructive cues and choice prompts were similar and significant, but decoding accuracy was generally lower for the latter; this was likely the result of there being only half as many trials in the choice condition.

### Attentional Modulation of Baseline Activity in Visual Cortex

Attention cueing improves task performance (Posner et al., 1980). The underlying neural mechanisms have been the subject of intense investigation. Kastner et al. (1999) cued subjects on a block-by-block basis to covertly attend the stimuli located in the upper right quadrant of the visual field or to passively view the stimuli. They reported an increase in BOLD signal during the anticipatory period after each block was cued but before targets were presented. The effect was observed in V1, V2, V4, temporal-occipital (TEO) regions (fusiform gyrus), and regions within the dorsal attention network (DAN) consisting of the frontal eye field (FEF) and intraparietal sulcus (IPS), with the BOLD activation being stronger in higher-order visual regions. Despite the seminal nature of this work, because the cued location was not varied in these studies in a fashion that could equate attention demands to and away from the attended region, it is difficult to know whether the patterns of the baseline activity encoded spatially-specific attended information or instead reflected nonspecific effects such as arousal or differences in the eccentricity of the focus of attention (i.e., fixation vs. peripheral). Hopfinger et al. (2000) contrasted the background biasing in visual cortex induced by attend-left versus attend-right cues—thereby equating the attention demands—and reported significant attention-related activations in higher-order visual areas in the fusiform and cuneus gyri (corresponding to V2, VP and V4) in the hemisphere contralateral to the attended hemifield, but not in V1. Sylvester et al. (Bressler et al., 2008; Sylvester et al., 2008; Sylvester et al., 2007; Sylvester et al., 2009), on the other hand, reported attention modulation in multiple visual areas including V1 during anticipatory attention. Our univariate analysis, combining both UF and UCD datasets through meta-analysis, showed baseline increases in BOLD activity during anticipatory attention in high-order visual areas such as lateral occipital and parietal regions, but not consistently in early retinotopic visual cortex, which is in agreement with the results of some prior studies (Hopfinger et al., 2000; O’Connor et al., 2002).

In voxel-based univariate analysis (e.g., whole-brain GLM analysis), for a voxel to be reported as activated by an experimental condition, it needs to be consistently activated across individuals. As such, individual differences in voxel activation patterns could lead to failure to detect the presence of neural activity in a given region of the brain. The univariate approach applied at an ROI level, on the other hand, may not have the spatial resolution to detect attention modulation that occurs at a sub-ROI level. For the latter, Silver et al. (2007) found that cue-related activity is enhanced in a subregion of V1 that corresponded to the attended stimulus, and more importantly, this enhanced region is surrounded by regions of deactivation, suggesting the formation of a neural pattern where there are both excitatory responses and inhibitory responses within the same ROI (Muller & Kleinschmidt, 2004; Smith et al., 2000). While such attention modulation patterns present a challenge for univariate analysis, they open the possibility for the fruitful application of multivariate pattern analysis (MVPA). Hypothesizing that attending to different spatial locations leads to different neural patterns, we applied the MVPA approach to the fMRI data during the cue-target period. Our results showed above-chance decoding in all retinotopic visual areas. The decoding accuracy strength was <u>higher in earlier visual areas</u> and decreased going up the visual hierarchy. This shows that top-down biasing signals influence baseline neural activity in early visual cortex during anticipatory spatial attention, and that the effect is not weaker than that in higher-order visual areas.

It is tempting to attribute our findings to the use of sensitive multivariate analysis methods, but multivariate analysis was also used by Park and Serences (2022). They also found differences in multivoxel patterns evoked by different attention-directing cues in several retinotopic extra-striate visual areas, but as in prior univariate studies, the effects during the anticipatory period were weaker in V1 than in extra-striate cortex. It is not clear why our decoding results and those of Park and Serences (2022) should differ, but we offer some thoughts on this. First, perhaps differences in the spatial scale of the target arrays was a factor. Hopf et al. (2006) argued that the spatial scale of the stimulus array (e.g., larger versus smaller) influences whether the selection occurs early (small scale) or late (large scale) in the visual hierarchy. Though both our study and the Park and Serences study used spatial gratings as the target stimuli, the stimulus array in Park and Serences (2022) was spread around the outline of an imaginary circle and the stimuli could occur at 12 possible locations, and this may have effectively increased the spatial scale of the stimulus array. In our study, the stimuli could appear at only two locations in the lower visual fields, and thus the spatial scale of our stimulus array was smaller. A smaller spatial scale would favor earlier selection, and this might explain why we observed robust decoding of attention conditions in V1. Second, during the cue-target interval in their design, Park and Serences presented a flickering noise stimulus; this bottom-up signal could have interfered with the nature of top-down activity in V1 (Uchiyama et al., 2021) and resulted in a weaker top-down signal. Future work will be required to resolve these differences between our work and that of Park and Serences.

### Behavioral Significance of Baseline Modulation in Visual Cortex

Do the biasing signals in the visual cortex impact behavior? This question has been addressed in previous research. Ress et al. (2000), using logistic regression modeling to relate relative BOLD activity in visual regions to behavior (the signal detection measure d’), observed the activity in V1-V3 to be predictive of behavioral performance. Sylvester et al. (2007) found the pre-stimulus neural activity in the visual cortex region v3A, indexed by subtracting the ipsilateral hemispherical BOLD activity (left hemisphere for attend-left; right hemisphere for attend-right) from contralateral BOLD activity (right hemisphere for attend-left; left hemisphere for attend-right), was higher for trials with correct responses than incorrect trials, again suggesting a relation between cue-evoked cortical activity and behavior. Stokes et al. (2009), applying a multivariate decoding approach to a cued attention experiment, found enhanced shape-specific baseline (pre-stimulus) neural activity in lateral occipital regions, and reported that both cue and stimulus-evoked decoding predicted target detection accuracy. However, in prior work from our group, Fannon et al. (2007) using a cued motion-color attention task, found that the increase in cue-related neural activity in the visual cortex (MT and V8) was not predictive of the post-stimulus neural activity or subsequent behavioral performance. Similarly, McMains et al. (2007) also reported differences in neural activity between baseline and target evoked activity. Thus, despite growing evidence of studies reporting associations between the behavior and baseline neural activity, the issue of whether cue-related neural activity impacts behavior remains an important issue to be better understood (Zheng et al., 2024).

We examined this issue using both univariate and MVPA approaches. We hypothesized that the differential activations from the univariate analysis and the decoding accuracy from MVPA (a measure of the strength of anticipatory attentional biasing) should be positively associated with behavioral performance. Our comparison of univariate activation difference (attend minus ignore) with behavioral efficiency (accuracy/RT), combining both UF and UCD datasets (through meta-analysis), failed to find a relation between the univariate BOLD activity in the retinotopic visual regions and behavioral performance. In contrast, the decoding accuracy from all the visual regions— whether as a whole as indexed by the first PCA component or analyzed on a ROI-by-ROI basis—were positively associated with behavioral efficiency, thus establishing a positive relationship between cue-related baseline activity in the visual cortex and subsequent behavioral performance.

### Study Design Considerations

Three aspects of our study design are worth emphasizing. The first is that we included two datasets recorded at two different institutions using two different brands of scanners but with the same experimental paradigm in our analysis. By doing so, we were able to combine the datasets via meta-analysis to enhance statistical power, thus addressing a concern that neuroimaging studies often are underpowered and suffer from low reproducibility (Bennett & Miller, 2010; Noble et al., 2017; Poldrack et al., 2017).

The second is in our experimental paradigm we included a choice cue in addition to standard instructional cues. Upon receiving the randomly occurring choice cue intermixed with instructional cues, the subject had to freely decide whether to attend the left or right visual field. While our paradigm was originally designed to investigate willed attention (Bengson et al., 2014; Rajan et al., 2019), the availability of the choice cue allowed us to rule out the possibility that the above-chance decoding of cue-evoked activity in early visual cortex simply reflected the processing of the different physical attributes of the different visual cues (Carlson & Wardle, 2015; Grootswagers et al., 2017), and thus to rule in that it reflected the differences in neural patterns underlying the two different attentional states (attend left vs. attend right). Importantly, the neural patterns between cue-left and cue-right and between choice-left and choice-right are highly similar, as indexed by the above-chance cross-decoding in which the classifiers trained on instructed trials were tested on choice trials, suggesting that the visual cortical biasing follows the same neural mechanisms irrespective of whether the attentional state arose from external instructions or from pure internal decisions. It is also worth noting, though, that because the number of choice trials were half that of the instructed trials, the effects from the choice trials were often weaker, whether it is the size of the decoding accuracy in the visual hierarchy or the correlation between decoding accuracy with behavior.

The third aspect of the study design bearing consideration is the stimulus array. As shown in Figure 1 and described in the Methods, small, single white dots (placeholders) marked the general location to which targets would be presented. While these were stationary (i.e., not flashing or moving), shifting attention to the dots might result in attention-related modulations of sensory signals from the contrast border represented by the dots, rather than baseline shifts of background neural activity in the complete absence of sensory signals. Microsaccades could also result in the dot markers serving as bottom-up sensory inputs due to the movement of the dot stimuli on the retina. Because the dots were present in both the attended and unattended visual hemifields, one might be tempted to assume that the physical sensory features of the display are equivalent for cue/choose-left versus cue/choose-right conditions. While this is true for the display independent of attention, it is not necessarily true for the well-known interaction of top-down spatial attention and bottom-up sensory signals. That is, one might predict a pattern of BOLD effects similar to what we observed if the activity was the attentional modulation of the dot stimulus itself. There are reasons, however, to believe the current findings and interpretations of baseline activity are valid. First, we saw no consistent increase in univariate baseline activity in V1d, which would have been expected if the lower visual field dots were in fact being modulated by attention. Second, the dots and the flashed gratings are quite different in their physical features, and it is the task-relevant grating stimuli that are the targets of attention in our design. Third, even if one were to assume that the dots were providing a bottom up signal that was differentially modulated by attention, and therefore could be confounded with pure baseline BOLD shifts in this design, that would not invalidate the basic finding here that the magnitude of top-down attention during the anticipatory period correlates with the later behavioral performance for task-relevant target stimuli. Further, our univariate BOLD results are generally consistent with findings from prior work that have (Hopfinger et al., 2000; Serences et al., 2004) and have not (Giesbrecht et al., 2006; McMains et al., 2007; Walsh et al., 2011; Weissman et al., 2002) included background markers of spatial locations where higher order visual regions contralateral to the attended location are more activated following the attention directing cue. Finally, our multivariate results differ from those of Park and Serences (2022) who also had highly salient placeholder stimuli (outline circles) throughout each trial, and thus it is <u>un</u>likely that our strong decoding of activity in lower order visual areas is simply due to the presence of placeholder stimuli because Park and Serences actually found weaker decoding in early visual areas, despite the presence of their placeholders.

## Summary and Conclusions

In this study, we used multivariate analysis of fMRI data recorded during a trial-by-trial visual spatial attention task to examine how top-down spatial attention control modulates anticipatory activity across the visual hierarchy. We found clear evidence for strong attentional biasing during the anticipatory period in all visual areas, including the primary visual cortex. Importantly, this effect was observed regardless of whether spatial attention was directed by external cues (instructed attention) or based on the participants’ internal decisions (choice attention).

These results contrast with previous studies that employed univariate methods, where baseline biasing during the anticipatory period was typically found to be stronger in higher-order visual areas, findings we also replicated here using univariate methods. One possible explanation for this discrepancy is that multivariate decoding methods are more sensitive, enabling the detection of neural activity patterns that might not be apparent with univariate approaches.

Importantly, we observed a significant positive relationship between decoding accuracy during the anticipatory period and subsequent behavioral performance on the target discrimination task. In other words, the stronger the decoding accuracy in the anticipatory period, the better participant’s performed in the task.

Overall, these findings provide new insights into how top-down attentional control biases visual processing across the visual hierarchy, and highlight the important relationship between anticipatory brain activity and human perception and performance.

## Acknowledgements

This work was supported by National Institutes of Health grant MH117991 and National Science Foundation grant BCS-2318886 to G.R.M. and M.D. We thank E. Ferrer, Y. Liu, J. Bengson, T. Kelley, K. Bo, Z. Hu, and S. Kim for discussion and help with data collection and analysis.

## Author Contribution

**Sreenivasan Meyyappan:** Conceptualization; formal analysis (lead); methodology (lead); writing – original draft (lead); writing – review and editing. **Abhijit Rajan**: Conceptualization; data curation; methodology; writing – original draft. **Qiang Yang**: data curation; formal analysis. **George R Mangun**: Conceptualization; funding acquisition; writing – review and editing. **Mingzhou Ding**: Conceptualization (lead); funding acquisition; writing – review and editing.

